# Production, purification, and kinetics of the poly-β-hydroxybutyrate depolymerase from *Microbacterium paraoxydans* RZS6: A novel biopolymer-degrading organism isolated from a dumping yard

**DOI:** 10.1101/540609

**Authors:** RZ Sayyed, SJ Wani, Abdullah A. Alyousef, Abdulaziz Alqasim, Asad Syed

## Abstract

Poly-β-hydroxybutyrate (PHB) depolymerase can decompose biodegradable polymers and therefore has great commercial significance in the bioplastic sector. However, few reports have described PHB depolymerases based on isolates obtained from plastic-contaminated sites that reflect the potential of the source organism. In this study, we evaluated *Microbacterium paraoxydans* RZS6 as a producer of extracellular PHB depolymerase isolated from a plastic-contaminated site in the municipal area of Shahada, Maharashtra, India, for the first time. The isolate was identified using the polyphasic approach, i.e., 16S rRNA gene sequencing, gas chromatographic analysis of fatty acid methyl esters, and BIOLOG identification, and was found to hydrolyze PHB on minimal salt medium containing PHB as the only source of carbon. Both isolates produced PHB depolymerase at 30°C within 2 days and at 45°C within 4 days. The enzyme was purified most efficiently using an octyl-sepharose CL*-*4B column, with the highest purification yield of 6.675 U/mg/mL. The enzyme required Ce^2+^ and Mg^2+^ ions but was inhibited by Fe^2+^ ions and mercaptoethanol. Moreover, enzyme kinetic analysis revealed that the enzyme was a metalloenzyme requiring Mg^2+^ ions, with optimum enzyme activity at 45°C (thermophilic) and under neutrophilic conditions (optimum pH = 7). The presence of Fe^2+^ ions (1 mM) and mercaptoethanol (1000 ppm) completely inhibited the enzyme activity. The molecular weight of the enzyme (40 kDa), as estimated by sodium dodecyl sulfate-polyacrylamide gel electrophoresis, closely resembled that of PHB depolymerase from *Aureobacterium saperdae*. Scale-up from the shake-flask level to a laboratory-scale bioreactor further enhanced the enzyme yield. Our findings highlighted the applicability of *M. paraoxydans* as a producer of extracellular PHB depolymerase isolated from a plastic-contaminated site in the municipal area of Shahada, Maharashtra, India.

## Introduction

Poly-β-hydroxy alkanoates (PHAs) and poly-β-hydroxybutyrate (PHB) are stored as food and energy sources in bacteria under a carbon-rich environment and are catabolized during nutrient stress conditions under the influence of PHB depolymerase. PHB is a biocompatible, thermoplastic, nontoxic, completely biodegradable molecule exhibiting the properties of synthetic plastics. Moreover, PHB is easily degraded by PHB depolymerases and hence is an eco-friendly alternative to recalcitrant synthetic plastics (1-5)(Sayyed and Chincholkar 2009; Gangure et al. 2017; Wani et al. 2016; Wani and Sayyed 2016; Soam et al. 2012).

PHB is degraded under natural conditions by the actions of PHB depolymerases produced by a wide variety of microorganisms (6, 7) (Papaneophytou et al. 2009; Hsu et al. 2012). Although PHB has commercial applications, identification of potent PHB degraders from relevant habitats and evaluation of their production, scale-up purification, and enzyme kinetic must be carried out. Reports on bacterial PHB depolymerases isolated from plastic-contaminated sites, which reflect the biodegradation potential of the enzyme, are scarce. Organisms isolated from plastic-contaminated sites that are capable of degrading PHB may serve as potential sources of efficient PHB depolymerases.

Accordingly, in this study, we aimed to isolate PHB-degrading bacteria from plastic-rich dumping yards. We describe the isolation and polyphasic identification of a PHB depolymerase-producing *Microbacterium paraoxydans* RZS6 isolated from plastic-contaminated sites and then the production, purification, characterization, and enzyme kinetics of the identified PHB depolymerase.

## Materials and Methods

### Chemicals and glassware

All the chemicals used in this study were of analytical research grade. PHB was purchased from Sigma-Aldrich (Germany); all other chemicals were purchased from Hi-Media Laboratories (Mumbai, India). The glassware was cleaned using 6N HCl and K_2_Cr_2_O_7_, rinsed with double-distilled water and dried in an oven.

### Isolation and screening of PHB depolymerase-producing bacteria

Sample collection, isolation, and screening of the PHB depolymerase-producing bacteria were performed as described by Wani et al. (2016) (3). *M. paraoxydans* RZS6 and *Stenotrophomonas maltophilia* RZS7 were isolated from plastic-contaminated sites located at latitude 21° 30’ 47.09″ N and longitude 74° 28’ 40.47″ E. These sampling sites were chosen purposefully due to the higher probability of finding microflora that would be metabolically very active in biopolymer degradation.

### Selection of potent isolates

Isolates producing the zone of PHB hydrolysis were grown on MSM containing different concentrations of PHB (0.1–0.4%) at 30°C for 10 days (8) (Pushpita et al. 2006). The degradation of PHB was detected by observing the time profile of the growth of the isolates, and by observing the formation of the zone of PHB clearance surrounding the colonies. The level of PHB degradation was measured from the diameter of the zone of PHB hydrolysis.

### Temperature profile of the potent isolates

In order to assess the thermostability of the PHB depolymerase, the isolate was subjected to PHB degradation assays for a period of 10 days at 28°C, 37°C, or 45°C in MSM containing different concentrations of PHB (0.1%, 0.2%, 0.3%, and 0.4%). The influence of temperature on PHB degradation was assessed by measuring the zone of PHB hydrolysis on each plate.

### Polyphasic identification of the isolates

Isolates showing the highest potential for PHB degradation on the PHB-agar were considered potent PHB depolymerase producers and were subjected to polyphasic identification.

#### Preliminary identification

Colonies of PHB-degrading isolates on nutrient agar (NA) medium were characterized using the Gram-staining method, morphological characteristic analysis, and taxonomic characterization by biochemical kits (Hi-Media, Mumbai, India). Isolates were identified according to Bergey’s Manual of Determinative Bacteriology (9) (Brenner et al. 2005).

#### 16S rRNA gene sequencing

Sequencing of 16S rRNA genes of the isolate RZS6 was performed as per the method of (10) Gangurde et al. (2013). DNA of the isolate was extracted according to the methods of (11) Sambrook and Russel (2001) using a HiPurA Plant Genomic DNA Miniprep purification spin kit. Amplification of the 16S rRNA genes was performed using the following primers: 27f (5’-AGAGTTTGATCCTGGCTCAG-3’) and 1492r (3’-ACGGCTACCTTGTTACGACTT-5’) (12) (Pediyar et al. 2002). The amplified sequences were analyzed using gapped BLASTn (http://www.ncbi.nlm.nih.gov), and the evolutionary relationship was computed using the neighbor-joining method in Clustal W software (13) (Thompson et al. 1997). Phylogenetic trees were constructed according to the methods described by Tamura and Kumar (2004) (14). The 16S rRNA gene sequences of the isolate were submitted to GenBank.

#### Fatty acid methyl ester (FAME) analysis

Fatty acids from whole cells of the isolate RZS6 derivatized to methyl esters were analyzed by gas chromatography using the Sherlock Microbial Identification System (MIDI, Inc., Newark, DE, USA) (15) (Gulati et al. 2008). Identification and quantification of fatty acids were carried out by comparing the retention time and peak area of the samples with those of standard fatty acids. Differences in the fatty acid profiles were computed using the Sherlock bacterial fatty acid ITSA1 aerobe reference library (16) (Sasser 2006).

#### Phenotypic fingerprinting

Phenotypic fingerprinting of isolate RZS6 was carried out using a GEN III MicroPlate test panel with 95 carbon source utilization assays (17) (Shaikh et al. 2014) and a Microbial Identification System, 1998 (Biolog Inc., CA, USA) with Micro Log version 4.2 database software (Biolog Microstation System; Biolog Inc.).

### Production and activity assay of PHB depolymerase

#### Growth kinetics and PHB depolymerase activity

Growth curve experiments were performed to evaluate the ability of the isolate to mineralize the substrate (PHB), as described by Maria and Zauscher (2002) (18). The growth rate of *M. paraoxydans* RZS6 in PHB MM was monitored over time at 620 nm in the presence of PHB as a substrate by withdrawing sample after every 12 h. The PHB depolymerase activity of the isolates was estimated as described by Papaneophytou et al. (2009) (6).

#### Production of PHB depolymerase

The production of PHB depolymerase was evaluated under shake-flask conditions by separately growing *M. paraoxydans* RZS6 at 30°C and 120 rpm for 4 days in MSM containing PHB (0.15%) (19) (Han and Kim 2002),

#### PHB depolymerase assay

After incubation, the MSM broth culture was centrifuged at 5000g for 15 min., and the PHB depolymerase activity of the supernatant was assayed as described by Papaneophytou et al. (2009) (6). *M. paraoxydans* RZS6 (5 × 10^6^ cells/mL) was grown in two reaction mixtures, each consisting of 50 mM Tris-HCl buffer (pH 7.0), 150 μg/mL PHB (prepared by sonication at 20 kHz for 15 min), and 0.5 mL of 2 mM CaCl_2_ at 30°C for 10 min. The PHB depolymerase activity was assayed as a decrease in the PHB turbidity at 650 nm. One unit of PHB depolymerase activity was defined as the quantity of enzyme required to cause a 0.1 decrease in absorbance at 650 nm per min.

### Purification of PHB depolymerase

Having confirmed the presence of a PHB depolymerase in the cell-free supernatant of the MSM broth culture, the supernatant was subjected to purification using three approaches, as described below.

#### Ammonium sulfate precipitation

The crude PHB depolymerase in the supernatant was precipitated by the gradual addition of increasing concentrations (10–40% w/v) of an ammonium salt. The obtained precipitate was dialyzed overnight (7) (Hsu et al. 2012), followed by estimation of the protein concentration and enzyme activity.

#### Solvent purification method

The culture supernatant of the isolate was centrifuged at 10,000 rpm for 20 min, and residues were dissolved in a pre-chilled 1:1 mixture of acetone and ethanol and kept in a water bath at 50°C to allow the evaporation of the solvent. The obtained pellet was dissolved in Tris-HCl buffer (pH 7), and the protein content and enzyme activity from the supernatant and pellet were assayed.

#### Column chromatography

The culture supernatant of *M. paraoxydans* RZS6 was loaded on to an octyl-sepharose CL*-*4B column charged with glycine*-*NaOH buffer (pH 9.0) and eluted using a 0–50% gradient of ethanol (20) (Kim et al. 2002). The fractions were collected and subjected to estimation of protein content and enzyme activity.

### Determination of molecular weight

The molecular weight of the purified PHB depolymerase of the isolate was determined using sodium dodecyl sulfate-polyacrylamide gel electrophoresis (SDS-PAGE) with standard molecular weight markers, such as phosphorylase B (82.2 kDa), bovine serum albumin (64.2 kDa), egg albumin (48.8 kDa), carbonic anhydrase (37.1 kDa), trypsin inhibitor (25.9 kDa), lysozyme (19.4 kDa), lysozyme (14.8 kDa), and lysozyme (6.0 kDa). The protein concentration of the purified band was measured using bovine serum albumin as a standard (21, 19) (Lowry et al. 1951; Han and Kim 2002).

### Enzyme kinetics

#### Effects of temperature on enzyme activity and determination of the thermostability of the enzyme

To determine the temperature optima and sensitivity of the PHB depolymerase, log culture of *M. paraoxydans* RZS6 (5 × 10^6^ cells/mL) was grown in the reaction mixture at different temperatures ranging from 5 to 70°C for 10 min, and the enzyme activity was then measured as described above.

#### Effects of pH on enzyme activity and determination of the pH stability of the enzyme

The effects of pH on enzyme activity and the pH stability of the enzyme were determined in reaction mixtures having varying pH values in the range of 2 to 13.

#### Effects of metal ions on the enzyme

In order to ascertain the metal requirement and type of metal required for the activity of the PHB depolymerase, *M. paraoxydans* RZS6 was separately grown in various reaction mixtures, each containing one type of metal ion, e.g., Ca^2+^, Mg^2+^, Mn^2+^, Cu^2+^, Co_2_^+^, Hg^2+^, Zn^2+^, and Fe^2+^ (1 mM), grown at 30°C for 10 min. Enzyme activity was then measured.

#### Effects of different chemicals on the enzyme

In order to determine the effects of solvents, namely, methanol (10%, v/v), ethanol (10%, v/v), acetone (10%, v/v), mercaptoethanol (1%, v/v), Tween*-*20 (1%, v/v), Tween*-*80 (1%, v/v), ethylenediaminetetraacetic acid (EDTA; 1 mM), NaCl (1 mM), KCl (1 mM), and NaNO_3_ (1 mM), on the activity of the PHB depolymerase. The solvents and chemicals were added individually into each reaction mixture, followed by inoculation with the isolate RZS6, incubation at 30°C for 10 min, and measurement of enzyme activity.

### Scale up of the optimized process to a laboratory-scale bioreactor

To evaluate the performance of the organism in the bioreactor and to confirm the validity of the optimized shake-flask studies, the process was scaled-up to a fully-automated bioreactor of 5-L capacity (Model LF*-*5; Murhopye Scientific Co., Mysore, India). The bioreactor was sterilized, along with the above-optimized medium (working volume 3 L), at 121°C for 20 min; the reactor was cooled and then inoculated individually with 3% (v/v) inoculum of *M. paraoxydans* RZS6. The samples were withdrawn after 12 h and subjected to estimation of protein concentrations (21) (Lowry et al. 1951) and enzyme activities.

## Statistical analysis

All the experiments were performed in triplicate and the mean of three replicates was considered. Each mean value was subjected to Student’s *t-*test and. Values of *P* ≤ 0.05 were taken as statistically significant (22) (Parker 1979).

## Results and Discussion

### Isolation and screening of PHB depolymerase-producing bacteria

In total, 39 isolates were obtained from the respective plastic-contaminated sites; among these 39 isolates, seven isolates grew well and produced varying degrees of the zone of PHB hydrolysis on minimal medium containing PHB as the only carbon source. The isolate RZS6 produced the largest zone of PHB hydrolysis (27.9 mm) and was therefore selected as the best PHB depolymerase producer. The hydrolysis of PHB reflected the ability of the isolate to produce PHB depolymerase.

Although several bacteria are known to secrete PHB depolymerase, which degrades PHB, our findings demonstrated, for the first time, the production of a PHB depolymerase from *M. paraoxydans* RZS6. The PHB depolymerase produced from these isolates (which were obtained from a plastic-contaminated environment) may be relevant for applications in plastic/bioplastic degradation. Mergaert et al. (1993) (23) also isolated 295 strains that degraded PHB and P (3HB-co*-*3HV) copolymer on MSM having PHB. Elbanna et al. (2004) (24) reported *Schlegelella thermodepolymerans* and *Pseudomonas indica* K2 as PHA degraders. Additionally, Gangurde et al. (2017) (2) also reported PHB biodegradation by soil micro-flora.

### Selection of potent isolates

PHB-degrading isolates exhibited a range of PHB biodegradation abilities in MSM containing varying concentrations of PHB. PHB degradation was dependent on the amount of PHB in the medium. Optimum biodegradation of PHB was recorded with 0.2% PHB (25) (Minna 2002). Augusta et al. (1993) (26) reported that the diffusion rate of the enzyme, level of enzyme activity, and incubation conditions affect the degradation rate. Moreover, Kim et al. (2003) (27) also reported similar observations with *Aspergillus* sp. strain NA*-*25.

### Temperature profile of the potent isolates

The isolate RZS6 exhibited optimum PHB degradation at 30°C with 0.2% PHB as a substrate. Importantly, the temperature profile of PHB degradation is dependent on the activity of the enzyme and therefore changes with the producing organism. Certain PHB depolymerases function in the mesophilic range of temperatures, whereas others are thermotolerant or thermophilic in nature (6) (Papaneophytou et al. 2009). Wang et al. (2012) (28) have reported similar observations on a poly-depolymerase (3-hydroxybutyrate-co*-*3-hydroxy valerate) from *Acidovorax* sp. HB01.

### Polyphasic identification of potent PHB depolymerase producers

#### Preliminary identification

The phenotypic characteristics of the potent PHB-degrading isolate RZS6 were similar to those of *Microbacterium* sp.

#### 16S rRNA gene sequencing

A BLAST search of the 16S rRNA gene sequences of the isolates with the 16S rRNA gene sequences of the NCBI GenBank database revealed the highest similarity and homology of the isolate RZS6 with *M. paraoxydans* (Fig 1). Thus, we identified the isolate as *M. paraoxydans*; the 16S rRNA gene sequence of the isolate was submitted to NCBI GenBank (http://www.ncbi.nlm.nib.gov/) under the name *M. paraoxydans* RZS6 (accession no. KP862607).

**Fig 1.**
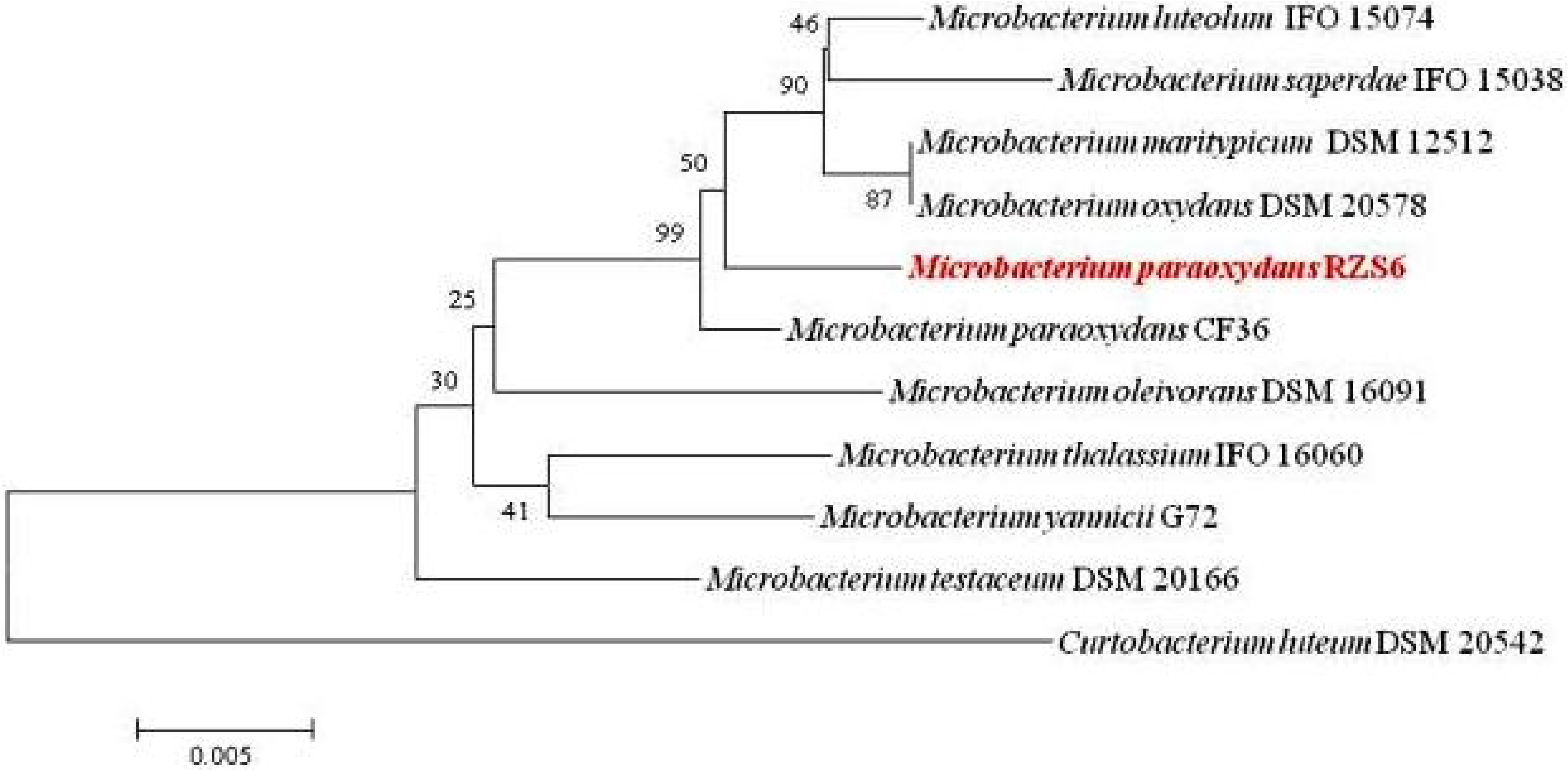
Phylogenetic analysis of *M. paraoxydans* RZS6 and related species by the neighbor-joining method using MEGA 5.0 software.

#### Whole-cell FAME analysis

The fatty acid profile of isolate RZS6 demonstrated the presence of characteristic fatty acids of *M. barkeri* (*Aureobacterium, Corynebacterium*; similarity index: 0.859) and *M. chocolatum* (similarity index: 0.602) (16) (Sasser 2006).

#### BIOLOG identification

The pattern of carbon source utilization assays for the isolate RZS6 demonstrated a maximum similarity index of 0.48 with *M. paraoxydans*. Based on the preliminary characteristics, 16S rRNA gene sequencing, gas chromatographic analysis of FAMEs, and BIOLOG profiles, the isolate RZS6 was identified as *M. paraoxydans*.

### Production and assay of PHB depolymerase

#### Growth kinetics and PHB depolymerase activity

Isolate RZS6 grew well in MSS containing PHB. The isolate showed faster growth, produced PHB depolymerase during the log phase of growth, and exhibited an optimum enzyme activity of 6.675 U/mg/mL, obtained at 48 h. The activity gradually decreased from the beginning of the stationary phase (72 h) and was completely absent during the decline phase (96–120 h) of growth (Fig 2).

**Fig 2.**
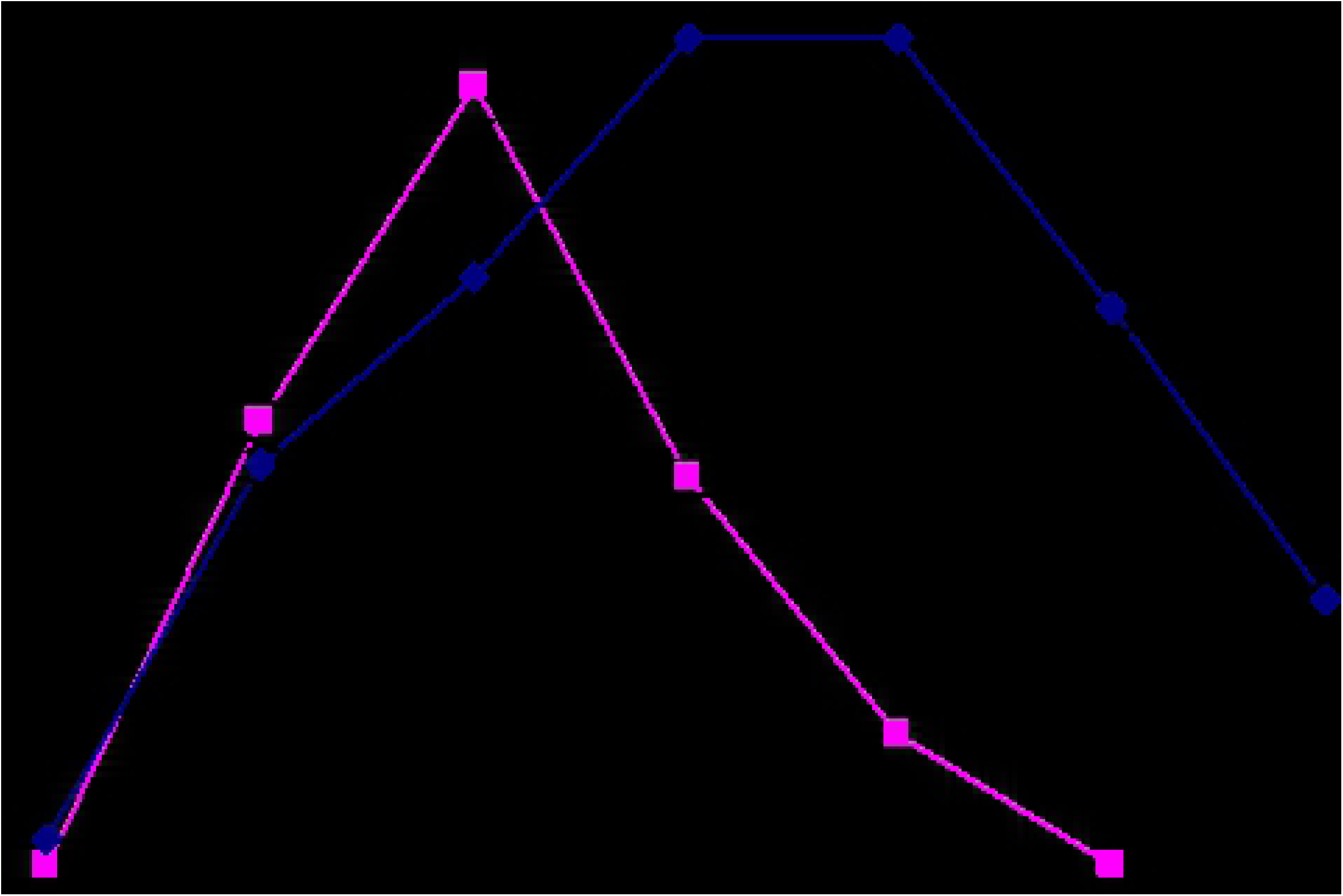
Growth kinetics and PHB depolymerase activity of *M. paraoxydans* RZS6.

#### Production of PHB depolymerase from the isolates

*M. paraoxydans* RZS6 produced copious amounts of PHB depolymerase in MSM. After 48 h (log phase) of incubation, RZS6 produced 6.6.75 U of PHB depolymerase with 0.247 mg/mL protein content in 2 days at 30°C. Gowda and Srividya (2015) (29) reported the production of 4U of extracellular PHB depolymerase with a protein content of 0.05 mg/mL from *Penicillium expansum*. Thus, our currently reported yields of the PHB depolymerase were higher than those of previously reported yields.

### Purification of the PHB depolymerase

#### Ammonium sulfate precipitation

The maximum protein precipitation in the culture supernatant was obtained at an ammonium sulfate concentration of 70%. The protein concentrations, specific activities, and enzyme activities in the dialyzed precipitate of *M. paraoxydans* RZS6 were 0.219 mg/mL, 0.321 mg/mL, and 6.6.75 U, respectively. Zhou et al. (2008) (30) also reported the precipitation of PHB depolymerase from *Escherichia coli* and *Penicillium* sp. DS9701-D2 using 70% and 75% ammonium sulfate. Shivakumar et al. (2011) (31) reported efficient precipitation of PHB depolymerase from *Penicillium citrinum* S2 using 80% ammonium sulfate.

#### Solvent purification method

Solvents adversely affected PHB depolymerase activity, with only approximately 45.66% and 51.14% remaining from *M. paraoxydans* RZS6. This significant loss in enzyme activity may be related to the precipitation of proteins (enzymes) by acetone and ethanol. Thus, the solvent purification method proved to be inefficient.

#### Column chromatography

Out of the five fractions, fraction 3 exhibited the maximum enzyme activity of 7.703 U, with 0.247 mg/mL protein content. Purification of PHB depolymerase from various organisms has been carried out using Sephadex columns, e.g., PHB depolymerase of *Bacillus* sp. (1.79 U/mg) (Shah et al. 2007) (32) *Streptoverticillium kashmirense* AF1, and *Streptomyces ascomycinicus* (Garcia et al. 2013) (33). Wang et al. (2012) (28) reported the purification of an extracellular PHB depolymerase from *Pseudomonas mendocina* DSWY0601 using a DEAE-Sepharose column. Papaneophytou et al. (2009) (6) and Hsu et al. (2012) (7) purified extracellular PHB depolymerases from *Thermus thermophilus* and *Streptomyces bangladeshensis* 77T*-*4 respectively using column chromatography. Thus, column chromatography using an octyl-sepharose CL*-*4B column resulted in the most efficient purification.

### Determination of the molecular weight of the purified enzyme

The purified protein fraction from *M. paraoxydans* RZS6 yielded single protein bands corresponding to molecular weights of approximately 40 kDa (Fig 3). Sadocco et al. (1997) (34) also reported a PHB depolymerase of 42.7 kDa from *Aureobacterium saperdae*. Calabia and Tokiwa (2006) (35) identified a PHB depolymerase of 41 kDa from *Streptomyces* sp. MG 41.

**Fig 3.**
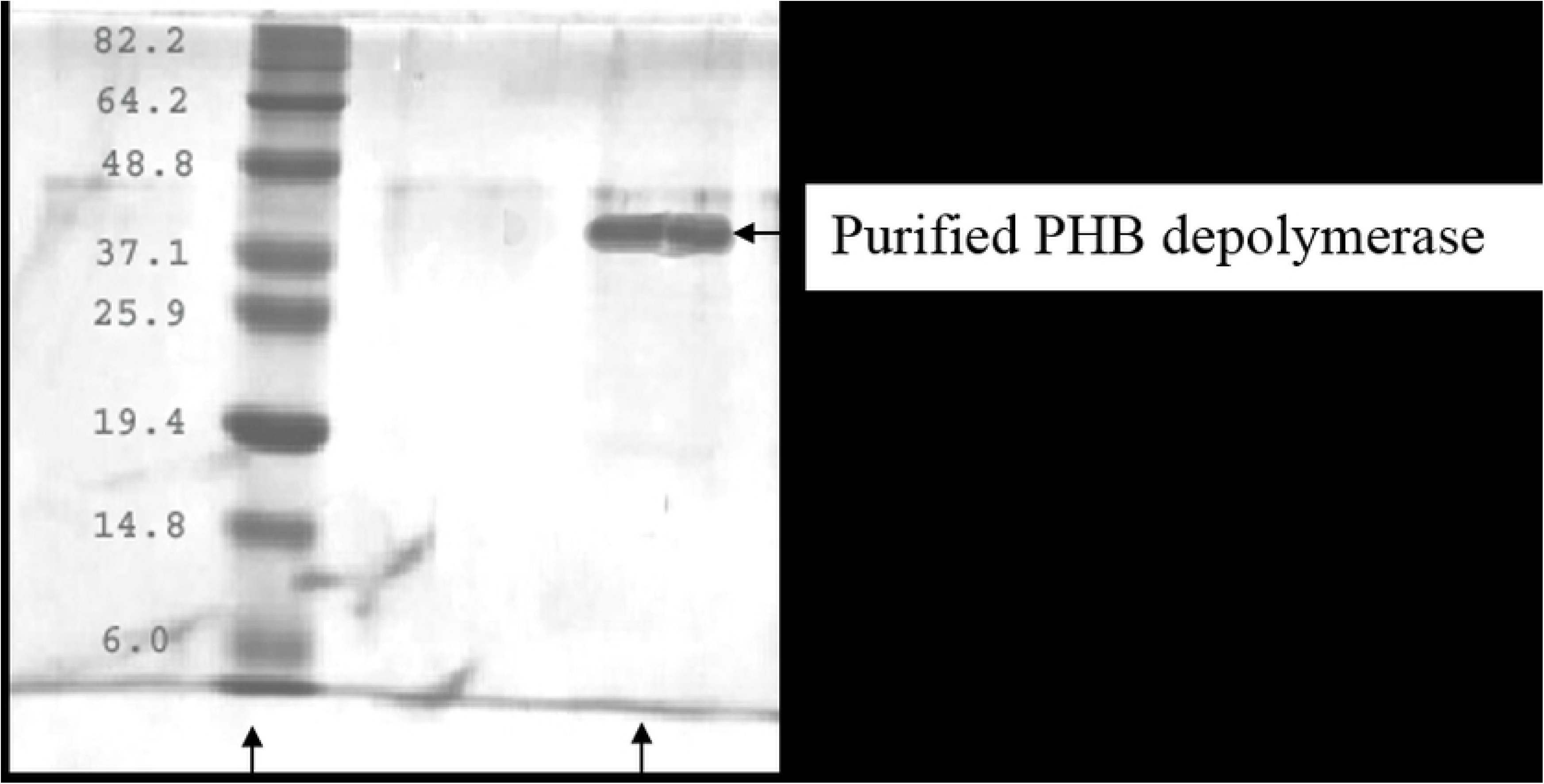
SDS-PAGE analysis of purified PHB depolymerase from *M. paraoxydans* RZS6.

### Enzyme kinetics

#### Effects of temperature and determination of the thermostability of the enzyme

The PHB depolymerase of RZS6 showed an optimum enzyme activity of 6.657 U with 0.247 mg/mL protein content at 30°C, indicating the mesophilic nature of the enzyme. The enzyme activity decreased as the temperature increased, and the enzyme was completely inactivated at 70°C (Fig 4). The decrease in the enzyme activity with the increase in temperature reflected the thermolabile nature of the enzyme. Wang et al. (2012) (28) and Gowda and Shivakumar (2015) (29) reported thermostable PHB depolymerases in *Pseudomonas mendocina* DSWY0601 and *Penicillium expansum*, respectively. Calabia and Tokiwa (2006) (35)reported optimum PHB depolymerase activity in *Streptomyces* sp. MG at 50°C.

**Fig 4.**
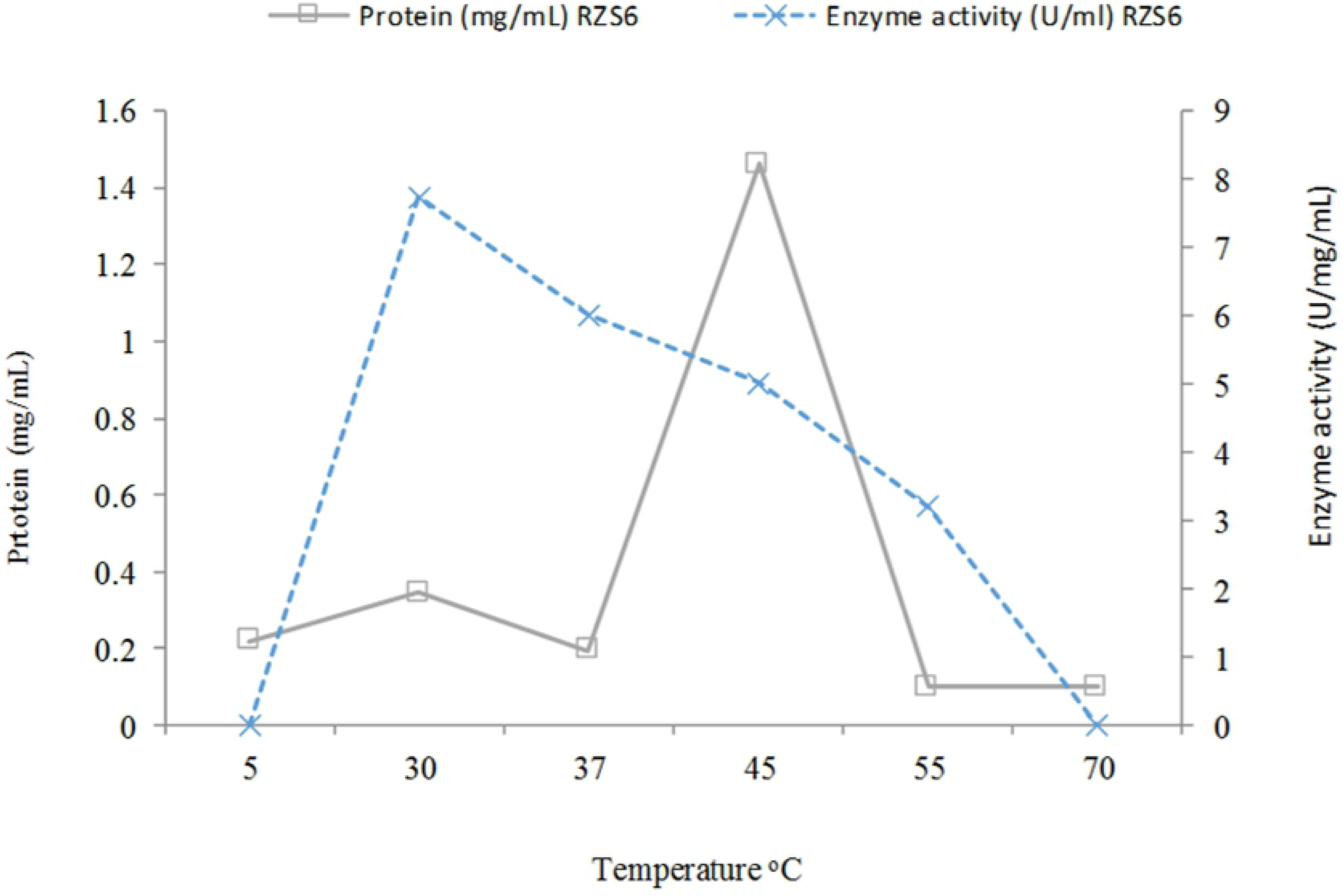
Effects of temperature on PHB depolymerase activity of *M. paraoxydans* RZS6.

#### Effects of pH on enzyme activity and determination of the pH stability of the enzyme

The optimum activity of the PHB depolymerase of *M. paraoxydans* RZS6 was obtained at pH 7 (Fig 5), indicating the neutrophilic nature of the enzyme. Sadocco et al. (1997) (34) and Jeong (1996) (36) reported an optimum pH for PHB depolymerase from *A. saperdae* in the range of 7.0 to 9.0. The pH sensitivity of the PHB depolymerases of *Penicillium expansum* and *Pseudomonas mendocina* DSWY0601 has been reported to be between 6.0 and 9.0 (Wang et al. 2012; Gowda and Srividya, 2015) (28,29). The enzymes from both isolates remained stable at an acidic pH (4.0–6.5). Soam et al. (2012) (5) reported pH 7.0 as the optimum pH for enzyme production in *Bacillus mycoides.* Calabia and Tokiwa (2006) (35) reported that the optimum PHB depolymerase activity of *Streptomyces* sp. MG was in the pH range from 6.5 to 8.5.

**Fig 5.**
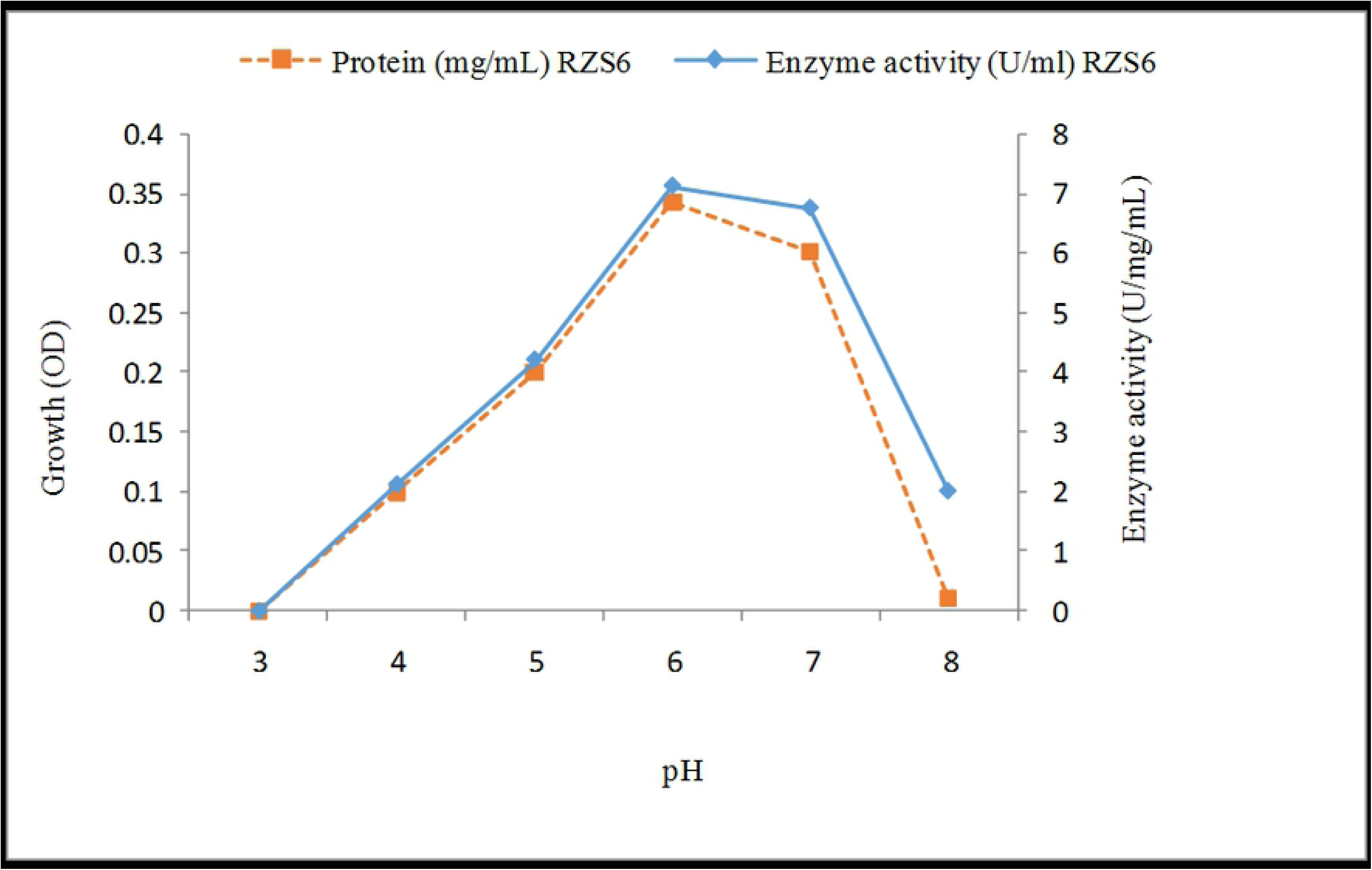
Effects of pH on PHB depolymerase activity of *M. paraoxydans* RZS6.

#### Effects of metal ions on enzyme activity

The presence of Mg^2+^ ions significantly enhanced PHB depolymerases. However, Fe^2+^ ions negatively affected the enzyme activity, decreasing activity to only 22.26%. The other metal ions had a negligible effect on the enzyme activities of both isolates (Table 1). The increase in PHB depolymerase activity in presence of Mg^2+^ ions was attributed to the enzyme activator nature of these metal ions. Wang et al. (2012) (28) also reported a positive effect of Mg^2+^ and Ca^2+^ ions on the PHB depolymerase of *Pseudomonas mendocina* DSWY0601.

**Table 1.** Effects of metal ions and chemicals on the PHB depolymerase of *M. paraoxydans* RZS6.

| **Metal ion**<br>**(1 mM)** | **Enzyme**<br>**activity (U/mL)** | **Protein content**<br>**(mg/mL)** | **Specific activity**<br>**(U/mg/mL)** | **%**<br>**activity** |
| --- | --- | --- | --- | --- |
| No metal ions | 0.146 | 0.247 | 0.6210 | **59.10** |
| $Mg^{2+}$ | 0.194 | 0.247 | 0.7854 | **78.54** |
| $Ca^{2+}$ | 0.192 | 0.247 | 0.7773 | 77.73 |
| $Mn^{2+}$ | 0.129 | 0.247 | 0.5627 | 56.27 |
| $Cu^{2+}$ | 0.099 | 0.247 | 0.4008 | 40.08 |
| $Co^{2+}$ | 0.101 | 0.247 | 0.4089 | 40.89 |
| $Hg^{2+}$ | 0.097 | 0.247 | 0.3927 | 39.27 |
| $Zn^{2+}$ | 0.094 | 0.247 | 0.3805 | 38.05 |
| $Fe^{2+}$ | 0.055 | 0.247 | 0.2226 | 22.26 |
| **Chemicals** |  |  |  |  |
| No chemical | 0.134 | 0.219 | 0.6118 | 61.18 |
| NaCl | 0.154 | 0.219 | 0.7196 | **71.96** |
| $NaNO_3$ | 0.148 | 0.219 | 0.6757 | 67.57 |
| Methanol | 0.145 | 0.219 | 0.6621 | 66.21 |
| KCl | 0.138 | 0.219 | 0.6301 | 63.01 |
| Ethanol | 0.112 | 0.219 | 0.5114 | 51.14 |
| Tween-80 | 0.11 | 0.219 | 0.5022 | 50.22 |
| Tween-20 | 0.103 | 0.219 | 0.4703 | 47.03 |
| Acetone | 0.091 | 0.219 | 0.4155 | 41.55 |
| EDTA | 0.046 | 0.219 | 0.2100 | 21.00 |
| Mercaptoethanol | 0.023 | 0.219 | 0.1050 | 10.50 |

#### Effects of different chemicals on enzyme activity

Mercaptoethanol caused the maximum inhibition (85%) of enzyme activity. NaCl enhanced the activity of the PHB depolymerase (Table 1). The loss of enzyme activity in the presence of chemicals and solvents occurs through denaturation and proteolysis of the enzyme. Papaneophytou et al. (2009) (6) also reported mercaptoethanol as a strong inhibitor of PHB depolymerase in *Thermus thermophilus* HB8. Wang et al. (2012) (28) have also reported the negative effects of ethanol, acetone, Tween, and other chemicals on the PHB depolymerase of *Pseudomonas mendocina* DSWY0601.

### Scale up of the optimized process to a laboratory-scale bioreactor

Scale-up of the parameters optimized at the shake-flask level to the laboratory-scale bioreactor level enhanced PHB depolymerase yield from 7.703 to 8.512 U and protein content from 0.247 to 0.297 mg/mL.

## Conclusion

Plastic-contaminated sites may harbor bioplastic (PHB)-degrading bacteria. The occurrence of PHB-degrading bacteria in plastic-contaminated sites and their growth in the presence of PHB indicated their ability to degrade bioplastic. Higher yields and optimal purification were obtained with column chromatography. The growth of the organism and the observation of maximum enzyme activity at higher temperatures are essential for the survival of the organism under extreme environmental conditions. Thus, PHB depolymerase-producing bacteria obtained from plastic-contaminated environments are expected to be important for applications in the biodegradation of bioplastics. However, further studies on the growth of *Microbacterium paraoxydans* and performance of PHB depolymerase under natural conditions shall help in this direction.

## Acknowledgments

The authors extend their appreciation to the Deanship of Scientific Research at King Saud University for funding this work through research group No (RG-1440-053).

## Author Contributions

**Conceptualization:** Christabel Ndahebwa Muhonja, Gabriel Magoma.

**Data curation:** Christabel Ndahebwa Muhonja, Huxley Makonde, Gabriel Magoma.

**Formal analysis:** Christabel Ndahebwa Muhonja, Huxley Makonde, Gabriel Magoma.

**Funding acquisition:** Gabriel Magoma.

**Investigation:** Christabel Ndahebwa Muhonja, Huxley Makonde, Gabriel Magoma, Mabel Imbuga.

**Methodology:** Christabel Ndahebwa Muhonja.

## Data Availability Statement

All relevant data are within the paper.

